# Censored Least Squares for Imputing Missing Values in PARAFAC Tensor Factorization

**DOI:** 10.1101/2024.07.05.602272

**Authors:** Ethan C. Hung, Enio Hodzic, Zhixin Cyrillus Tan, Aaron S. Meyer

## Abstract

Tensor factorization is a dimensionality reduction method applied to multidimensional arrays. These methods are useful for identifying patterns within a variety of biomedical datasets due to their ability to preserve the organizational structure of experiments and therefore aid in generating meaningful insights. However, missing data in the datasets being analyzed can impose challenges. Tensor factorization can be performed with some level of missing data and reconstruct a complete tensor. However, while tensor methods may impute these missing values, the choice of fitting algorithm may influence the fidelity of these imputations. Previous approaches, based on alternating least squares with prefilled values or direct optimization, suffer from introduced bias or slow computational performance. In this study, we propose that censored least squares can better handle missing values with data structured in tensor form. We ran censored least squares on four different biological datasets and compared its performance against alternating least squares with prefilled values and direct optimization. We used the error of imputation and the ability to infer masked values to benchmark their missing data performance. Censored least squares appeared best suited for the analysis of high-dimensional biological data by accuracy and convergence metrics across several studies.

## 1. Introduction

Dimensionality reduction methods summarize multivariate data by constructing a smaller set of explanatory features. Classical examples of these tools for data reduction, visualization, and unsupervised exploratory studies include principal component analysis and nonnegative matrix factorization. Due to the size of data and complex correlations among these variables in biological research, dimensionality reduction methods have widespread application^1,2^. Most methods use data structured in a tabular/matrix format, but measurements across several degrees of freedom, or modes (e.g., subjects, perturbations, time points, etc.) may be better represented as higher-dimensional arrays, or tensors^3^. Organizing data in a tensor form may better represent the experimental structure of these datasets, preserving patterns that might otherwise be lost when flattening data into matrix form to apply classical techniques.

Canonical polyadic decomposition (CPD), a natural extension of matrix factorization, is a common dimensionality reduction approach for analyzing data organized in tensor form^4^. It aims to represent the original tensor as the sum of several simpler tensors, each represented as the outer product of vectors associated with each mode. When appropriate, CPD decomposes data into a series of interpretable patterns describing variation across each mode. Tensor factorization, including CPD, can permit more extensive data reduction over matrix methods while also better elucidating patterns in co-dependent processes^5^. Previous work has illustrated the application of CPD in gene expression^6^, serum antibody analysis^7^, immune cytokine responses^8^, and various other fields^9^.

Missing values are a common and substantial hurdle for biological data analysis^3,10–12^. Missing values may arise from several sources, including discrepancies in which measurements were collected when integrating data from heterogeneous sources, limitations in experimental design, and data quality^13^. Missing values can be located in differing patterns across a dataset, such as a small number of individual values across the data^11^, or a more structured pattern such as all measurements of a feature across many samples. A variety of imputation methods have been developed to predict missing values from existing measurements, each assuming different existing relationships within the data^14–16^. For example, factorization-based imputation assumes the original data can be approximated by lower-rank matrices and seeks to fit the optimal factors from existing data^17,18^.

Imputation may additionally be used to assess the validity of a factorization^19^. To benchmark the ability to impute values, one can mask a subset of the data before applying tensor decomposition to impute these values, thus utilizing imputation as a form of cross-validation. Minimal differences between the masked and imputed values support that the method models observed measurements with high fidelity, thus providing support that biological patterns found within the factors are robust^20^.

Tensor decomposition presents both challenges and opportunities for handling missing values. Transforming a previously matrix-structured dataset into a tensor can itself introduce new missing values if not all combinations of features are measured. On the other hand, the tensor structure aligns measurements across shared modes, providing additional context with which to predict each entry^21,22^. Tensors reconstructed from solved factors are always complete, and so reconstructed values provide imputation predictions^4^. CPD can be solved by several algorithms, and the choice of algorithm affects the resulting factors. All approaches are reliant on an initial guess for the factor matrices, typically random values or the truncated singular value decomposition (SVD)^23^. One fitting approach is gradient-based direct optimization (DO), where all factors are solved simultaneously by treating all factors’ values as unknown parameters to be optimized by standard approaches^24^. DO can simply ignore missing values in the original tensor by removing these entries from the error function. However, this method suffers from numerical instability and difficulty in scaling to much larger and higher-dimensional tensors^25,26^. By contrast, alternating least squares (ALS) iteratively solves one mode at a time via linear least squares while holding the other modes constant^4^. ALS requires that missing entries are filled with values; one can repeatedly update these entries with the reconstructed values from the new decomposition in a process known as ALS with single imputation (ALS-SI)^26^. However, the prefilled values inevitably affect the least squares solution, introducing bias to the process. Neither method is inherently designed to address the factorization of datasets involving missing values.

Here, we propose censored alternating least squares (C-ALS) in place of ALS-SI in the factorization procedure when dealing with missing data. C-ALS harnesses the column independence of linear least squares and only considers existing values during factor solving, leveraging the computational stability of the ALS algorithm while preventing bias from prefilled values. To demonstrate the efficacy of C-ALS, we compared it against ALS-SI and DO on imputation accuracy and convergence time when running on four distinct biological datasets. We benchmarked each algorithm across several missing value patterns and amounts, finding that C-ALS generally yielded the best overall imputation compared to DO and ALS-SI. These tests also characterize the behavior of each algorithm, providing a systematic approach for method selection in the presence of missing values.

## 2. Methods

### 2.1 Notation

In this work, we denote scalars in italic (e.g. *a* or *A*), vectors in lowercase boldface (e.g. ***a***), matrices in boldface uppercase (e.g. **A**), and n-dimensional tensors with at least 3 modes in script typeface (e.g. ***𝒜***).

### 2.2 Introduction to the CP Decomposition

Tensors are multidimensional arrays. The number of modes indicates their dimensions. For example, a three-mode tensor, which is a three-dimensional array in the shape of *I* × *J* × *k* can be annotated as ***𝒯*** ∈ ℝ ^*I×J×K*^. Through CP decomposition, tensor ***𝒯*** can be approximated as the sum of several rank-one tensors and expressed as^27,28^

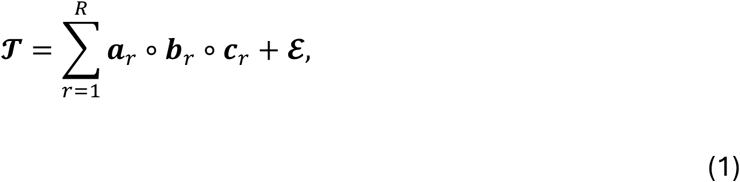

where *R* is the number of components, and ***a***_*r*_, ***b***_*r*_, and ***c***_*r*_ are the *r*-th factor for each of the three modes. These factors across *R* components can also be arranged into matrices, where ***a***_*r*_, ***b***_*r*_, and ***c***_*r*_ are the *r*-th columns of factor matrices **A** ∈ ℝ ^*I×R*^, **B** ∈ ℝ^*J×R*^, and **C** ∈ ℝ ^*K×R*^, respectively. “∘” represents the outer product, and ℰ is the residual that cannot be explained by these decomposed factors. Using the factor matrices, the original expression is equivalent to

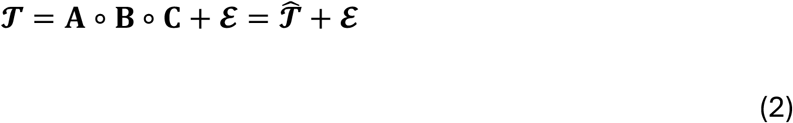

Solving the CP decomposition is equivalent to an optimization problem that tries to fit the reconstructed tensor, 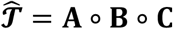, as close to the original tensor, ***𝒯***, as possible. It aims to minimize the following loss function with respect to the factor matrices, **A, B**, and **C**:

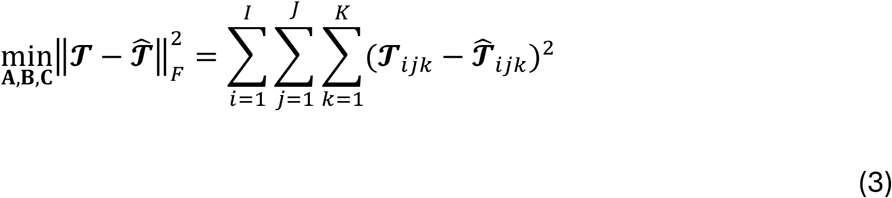

### 2.3 Solving CP decomposition with ALS

A common algorithm for solving the CP decomposition is ALS^18^. Essentially, ALS solves the factor matrix of one mode at a time with linear least squares while fixing the other factor matrices, then it repeats this process across all modes until the factors converge^28,29^. Here, we briefly demonstrate this method for a three-mode tensor.

First, we define the unfolding of tensor ***𝒯*** ∈ ℝ^*I×J×K*^ on its first mode as

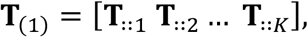

where **T**_∷*k*_ represents the matrix formed by {*T*_*ijk*_}_*i* = 1,…,*I*; *j* = 1,…,*j*_, the *k*-th frontal slice of tensor ***𝒯***, and the bracket “[ ]” concatenates these matrices into one. Therefore, an unfolding is a matricization of its original tensor format, and the unfolding of tensor ***𝒯*** in its first mode, **T**_(1)_, has dimension *I* × *Jk*. Similarly, the unfoldings of ***𝒯*** on the other two modes are **T**_(2)_ ∈ ℝ ^*J*×*IK*^ and **T**_(3)_ ∈ ℝ ^*K*×*IJ*^.

We define the Khatri-Rao product of matrices **B** ∈ ℝ^*J×R*^ and **C** ∈ ℝ ^*K×R*^ as

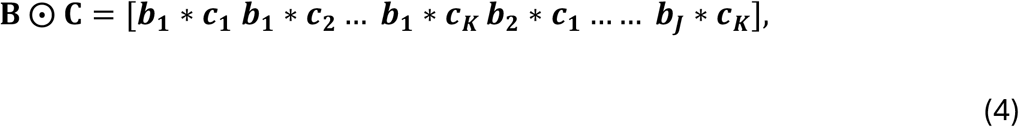

where ***b***_***j***_ ∈ ℝ^*R*^ is the *j*-th row of **B**, and ***c***_***k***_ ∈ ℝ^*R*^ the *k*-th row of **C**. “∗” indicates the elementwise product, and the bracket “[ ]” concatenates these vectors into a matrix. By this setting, the resulting product, **B ⨀ C**, has dimension *Jk* × *R*.

With these definitions, the minimization problem of 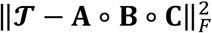 on **A** while holding **B** and **C** constant is equivalent to

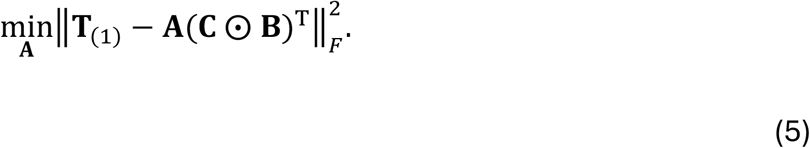

The optimal solution of **A** is given by linear least squares as

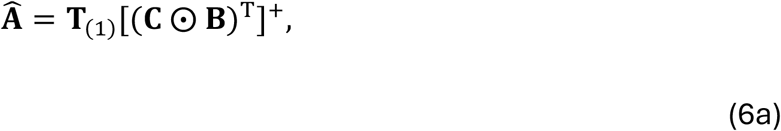

where “+” indicates the pseudoinverse of a matrix. Similarly, 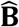 and 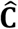 can be solved, respectively, as

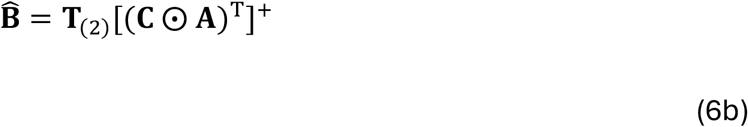

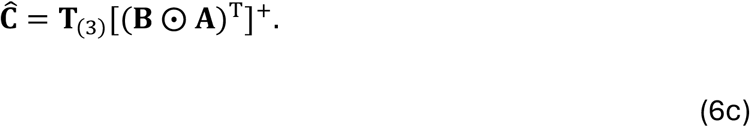

ALS first randomly initializes the factor matrices, **A, B**, and **C**, then iteratively updates them by Eqs. (6a)–(6c) until any changes are smaller than a given threshold.

### 2.4 Censored Alternating Least Squares (C-ALS)

When the original data tensor ***𝒯*** contains missing values, all its unfoldings, **T**_(1)_, **T**_(2)_, …, will contain missing values at corresponding positions as well. Therefore, the solution given by Eq. (6) is no longer viable. C-ALS replaces the step of Eq. (5) with a censored version of linear least squares, aiming to solve Eq. (5) with missing values in **T**_(1)_. Here, **W** = (**C ⨀ B**)^T^ ∈ ℝ ^*R×JK*^ is a complete matrix, and **T**_(1)_ ∈ ℝ ^*I×JK*^ is a matrix with missing values. The missing values in matrix **T**_(1)_ can be encoded by a masking matrix, **M**_(1)_ ∈ {0, 1} ^*I×JK*^, where (**M**_(1)_)_*ip*_ = 0 if (**T**_(1)_)_*ip*_ is missing, and (**M**_(1)_)_*ip*_ = 1 otherwise. The original loss function, Eq. (5), now becomes

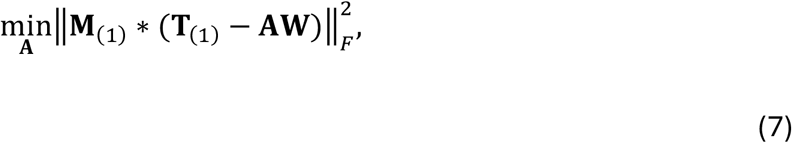

and the solution of **A** ∈ ℝ^*I×R*^ contains no missing values.

**A** can be solved by each row. The *i*-th row of 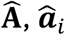, is given by

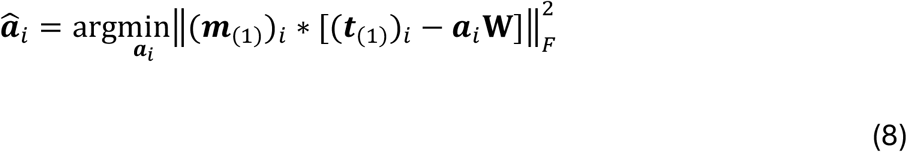

where (***t***_(1)_)_*i*_ is the *i*-th row of **T**_(1)_, and (***m***_(1)_)_*i*_ is the *i*-th row of **M**_(1)_.

Censored least squares proposes that when there are missing values in (***t***_(1)_)_*i*_, ***a***_*i*_ can be solved by performing linear least squares only on the submatrix of **W** where (***t***_(1)_)_*i*_ has values, ignoring the missed rows. Specifically, its solution for ***a***_*i*_ is

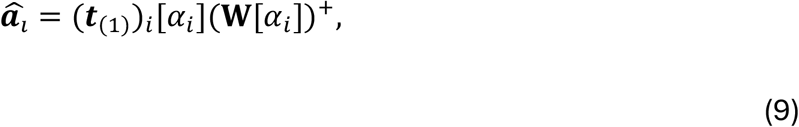

where *α*_*i*_ = {*p*∣(**M**_(1)_)_*ip*_ = 1} ⊆ {1,2, …, *Jk*} is the set of all indices where values in (***t***_(1)_)_*i*_ are not missed such that the cardinality of *α*_*i*_, |*α*_*i*_|, should be exactly the number of non-missing entries in 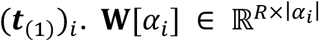 is a submatrix of **W** that only contains columns with indices in set *α*_*i*_, and 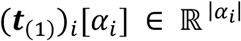 is a subvector of (***t***_(1)_)_*i*_. In other words, (***t***_(1)_)_*i*_ [*α*_*i*_] is exactly (***t***_(1)_)_*i*_ after leaving out all missing values, and **W**[*α*_*i*_] leaves out the corresponding columns in **W** as well. In the context of solving tensor decomposition with missing values, C-ALS replaces Eqs. (6a)–(6c) with

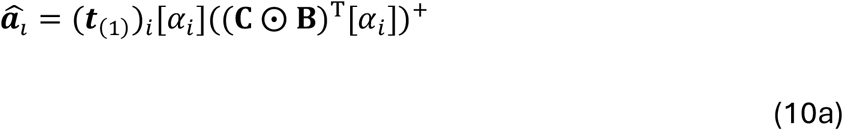

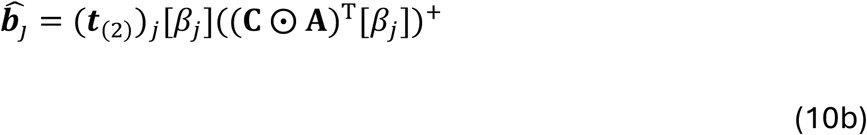

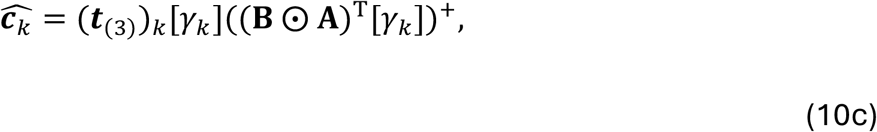

where *β*_*j*_ = {*p*∣(**M**_(2)_)_*jp*_ = 1} ⊆ {1,2, …, *Ik*} and *γ*_*k*_ = {*p*∣(**M**_(3)_)_*kp*_ = 1} ⊆ {1,2, …, *IJ*}, and **M**_(2)_ and **M**_(3)_ are the masking matrices of **T**_(2)_ and **T**_(3)_, respectively. The rest of the alternating least squares scheme remains the same.

### 2.5 Other methods for comparison

To benchmark the performance of C-ALS, we implemented two widely used methods missing data tensor factorization, ALS with single imputation and direct optimization.

#### 2.5.1 Alternating Least Squares with Single Imputation (ALS-SI)

The ALS-SI method follows an iterative approach to the expectation-maximization scheme^26^. Based on the ALS algorithm, missing values are first imputed via the initialization step, and the factor matrices are then iteratively updated until convergence. Let the randomly initialized factor matrices be **A**^(0)^, **B**^(0)^, and **C**^(0)^. The reconstructed tensor from these factors is

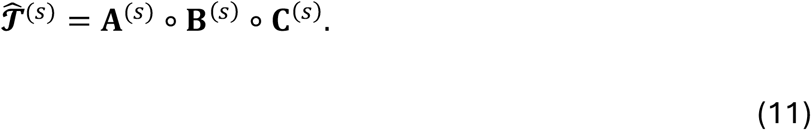

At the beginning, 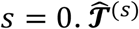 contains no missing values. ALS-SI constructs an imputed tensor, 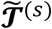, where any missing values in the original tensor, ***𝒯***, will be filled with their corresponding values in 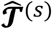:

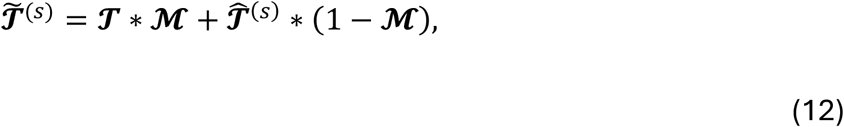

where ℳ ∈ ℝ^*I×J×K*^ is the masking tensor of ***𝒯*** (ℳ_*ijk*_ = 0 iff ***𝒯***_*ijk*_ is missing, otherwise ℳ_*ijk*_ = 1). As 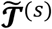 contains no missing values, ALS can be performed by unfolding it on each mode and the factor matrices are updated according to Eqs. (6a)–(6c). The resulting factor matrices, **A**^(s+1)^, **B**^(s+1)^, and **C**^(s+1)^ will be plugged into Eq. (11) to again to update the reconstructed tensors, 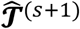. This process is repeated until the factors converge.

#### 2.5.2 Direct Optimization (DO)

Since CP factorization is, in its essence, an optimization problem (Eq. 4), it can be solved using a gradient-based optimization approach^30^. Considering the missing values in ***𝒯***, we have

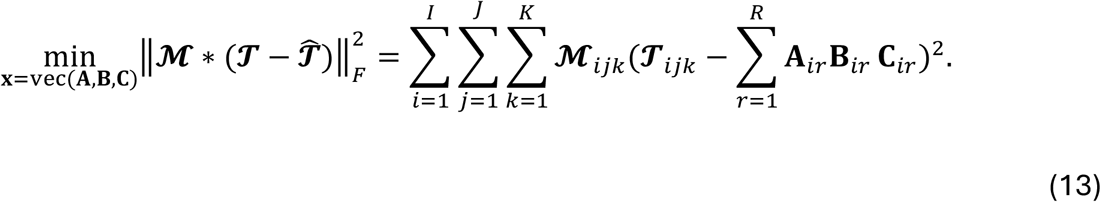

Here, the reconstructed values at the missed position are not included in the loss function. In this study, we used the limited memory Broyden–Fletcher–Goldfarb–Shanno algorithm with bound constraints (L-BFGS-B) implemented by SciPy^31–33^. In practice, the values in the factor matrices are flattened into one vector. After random initialization for the factor matrices, steps are taken until a maximum of 50 iterations or the function change was smaller than tolerance, 10^−6^.

## 3. Experiments

### 3.1 Data

We compare these decomposition algorithms with four datasets:

- **SARS-COV-2 serology** Zohar *et al*, 2020 profiled SARS-CoV-2-negative and - infected patients over about 4 weeks using systems serology^34^. Antibodies were profiled for their antigen and Fc receptor engagement. The data was organized into a tensor of size 432 × 6 × 11 corresponding to the sample, antigen, and receptor measured with no pre-existing missing values.
- **HIV serology** Alter *et al*, 2018, provides a second systems serology dataset collected from HIV-infected patients with distinct disease control, including elite controllers, treated progressors, untreated progressors, and viremic controllers^35^. Among other measurements, antibodies were profiled for their antigen and Fc receptor engagement. The data was organized into a tensor of size 181 × 41 × 22 corresponding to the sample, antigen, and receptor measured. This structure leads to 43% pre-existing missing values.
- **DyeDrop profiling** Mills *et al*, 2022, provides a high-throughput, single-cell, dose-response dataset profiling the cytotoxic and cytostatic properties of investigational pharmaceutical agents on a variety of breast cancer (BC) cell lines using a public domain Dye Drop method^36^. Cell lines were treated with agents at up to (but not necessarily always) 10 dosage levels and two time points. Cell viability and DNA replication assays were used to identify cell phase and subsequently followed by immunofluorescence imaging (per the Dye Drop protocol). Cell lines with more than 50% missing data and agents evaluated with six or fewer doses were left out, leaving data organized into a tensor of size 280 × 8 × 54 corresponding to agent at a given dosage, cell cycle phase at each of two timepoints, and cell line. 2% of the values were missing.
- **Breast cancer (BC) cytokine** Orcutt-Jahns *et al*, 2023, characterized peripheral blood mononuclear cells (PBMCs) in response to various cytokine treatments^37^. The phosphorylation responses of transcription factors were measured for various cell populations, leaving data that was organized into a tensor of size 36 × 8 × 23 × 6, corresponding to each sample, treatment, cell population, and cell marker, with 10% pre-existing missing values. Note that for random chord-wise masking, chords were left out along the sample mode.

For all experiments, factorizations were run on 20 randomly generated patterns of the indicated structure (see 3.2), with random initialization using unique seeds (generating unique initializations for each run of the method) that were reused for each method.

### 3.2 Imputation Masking

To evaluate the imputation capability of these methods, we hid a portion of the non-missing data from each method, imputed these data, and then compared the imputed and true masked values. The resulting error of was defined as:

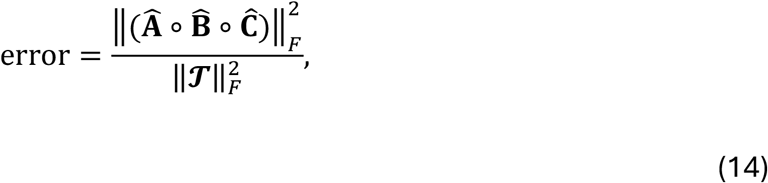

where the imputation error is calculated only on masked values and fitting error is calculated only on unmasked values. Pre-existing missing values were ignored.

The masked values were sampled in two separate fashions: entry and chord. When values were masked by random removing entries, a random subset of all non-missing positions was selected. Alternatively, when values were removed by chord (or fiber), all values were dropped along a select mode for a tensor after selecting random subset among indices of the remaining modes. For example, for ***𝒯*** ∈ ℝ ^*I×J×K*^, a chord along its first mode can be ***t***_:*jk*_, a vector formed by {*T*_*ijk*_}_*i* = 1,…,*I*_. We randomly chose among all possible chords for masking.

### 3.3 Performance Metrics/Rubrics

To determine the number of components for the decomposition, we solved for the CP model using 1 to 10 components for 3-mode datasets and up to 20 components for the 4-mode dataset. Where applicable, the factorization with the lowest median imputed error is selected, otherwise opting for the lowest fitting error in factorizations without imputation.

To evaluate the performance of these imputation methods, we ran the factorization multiple times with different starting points and plotted the medians and interquartile ranges of the errors from the runs. All methods were set to terminate at the same tolerance, a relative change of 10^-6^. When comparing method runtimes, median errors and time for each iteration were used to provide a demonstration of algorithm behavior.

## 4. Results

### 4.1 CP Decompositions Permit Tensor Analysis and Imputation

While standard matrix factorization approaches require flattening data across modes until a second-order tensor is achieved, construction of dataset into a higher-order tensor format permits the data to be expressed in a more native, combinatorial structure (Fig. 1a). CPD aims to generate component factors that maximally explain a dataset’s variation (Fig. 1b). These factor matrices may also be used to directly gain insight into the underlying factors or reconstruct the original tensor to evaluate the fidelity of the factorization. This reconstructed tensor has no missing values and thus the factorization may be treated as an imputation process as well. Several approaches exist for generating a CPD factorization for any given tensor. C-ALS is introduced here with the expectation that solving similarly missing portions of the data in separate least squares problems may increase the resource and time consumption of the solving process but improve fitting performance. C-ALS is thus compared to the ALS-SI and DO to benchmark the capabilities of the C-ALS algorithm (Fig. 1c). To assess each algorithm, an imputation scheme is devised whereby a percentage of randomly selected values in the original dataset is artificially masked in either an entrywise or chordwise before computing the CPD (Fig. 1d). The tensor reconstructed from the factor matrix is subsequently evaluated for masked (imputation) and unmasked (fitting) error for the CPD of each dataset at various component ranks.

**Fig. 1.**
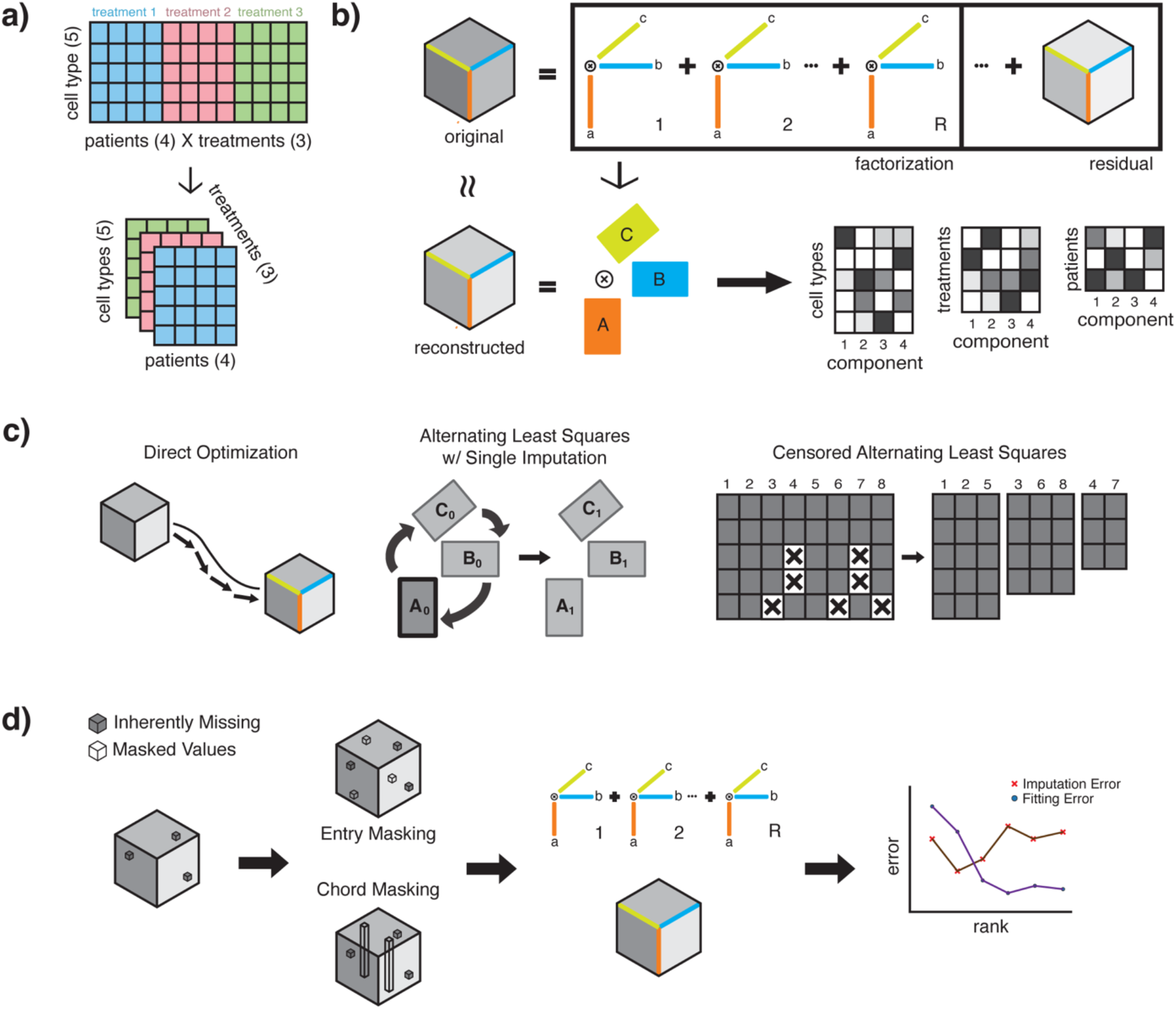
Tensor factorization and imputation approaches. **(a)** In multimodal datasets with three or more dimensions, tensor methods improve upon matrix methods that would otherwise concatenate combinations of factors, thus forfeiting relevant patterns across all modes. **(b)** With canonical polyadic decomposition (CPD), an original tensor is decomposed into R components, with each mode represented by a vector. The factor matrices may also be reconstructed into a new tensor that approximates the original. **(c)** A visual comparison of each algorithm: DO performs gradient descent with the vectorized tensor, ALS-SI iteratively alternates solving the least square solution for each mode after initial imputation of missing values, and C-ALS groups vectors at each linear least squares problem by their missing value patterns before proceeding to ALS for each group separately. **(d)** To evaluate imputative capabilities, values are artificially masked (on top pre-existing missing values) before running the method and assessed for reconstructive fidelity of withheld (imputation) and observed (fitting) values.

### 4.2 Factorization Fit before Imputation

To provide baseline performance for the three factorization algorithms, we first evaluated their fitting performance on the four datasets without masking (Fig. 2). In all four cases, C-ALS slightly outperformed DO. In the SARS-COV-2 serology dataset, ALS-SI and C-ALS performed identically; the original data contain no missing values and so these two algorithms are identical. The DyeDrop profiling which contains few missing values also displays similar patterns. In the HIV serology and BC cytokine datasets, C-ALS outperformed ALS-SI at higher component ranks.

**Fig. 2.**
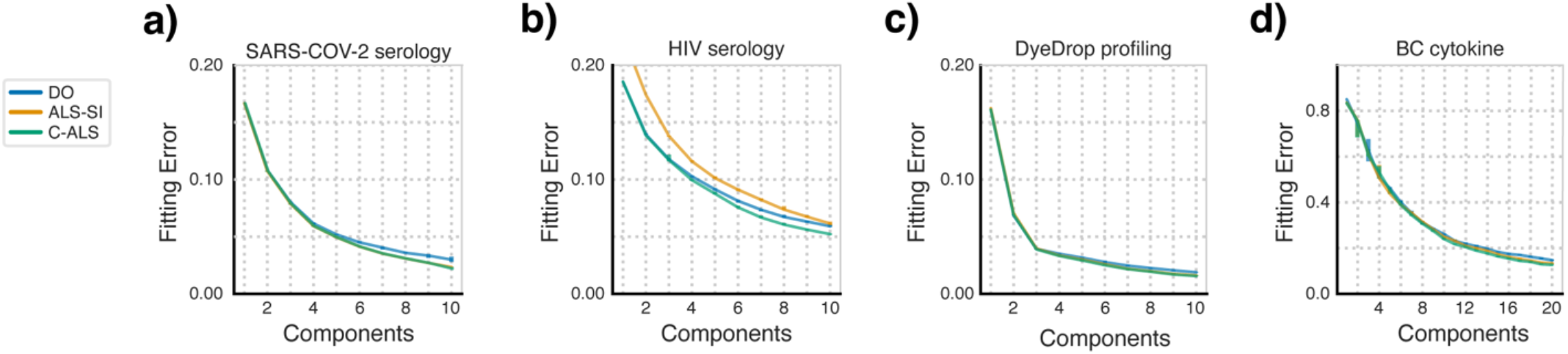
Least squares methods outperform gradient descent in fitting error. The fitting error of the three factorization algorithms with no masked values is shown. Data points represent the median error with error bars representing the inter-quartile range. N=20 for all tests.

### 4.3 Factorization Imputation

To compare the performance of imputing missing entries within a dataset, we ran the three factorization algorithms on all datasets with 10% of the entries masked (Fig. 3a–d). Across the datasets, C-ALS outperformed the other methods in both fitted and imputed error. The imputed error decreased monotonically with rank, except for the DyeDrop profiling dataset, where ALS-SI only improved up to 7 components before gradually increasing in imputed and fitted error (Fig. 3c). As with the non-missing case, C-ALS outperforms ALS-SI in terms of fitting and imputation error at higher numbers of components.

**Fig. 3.**
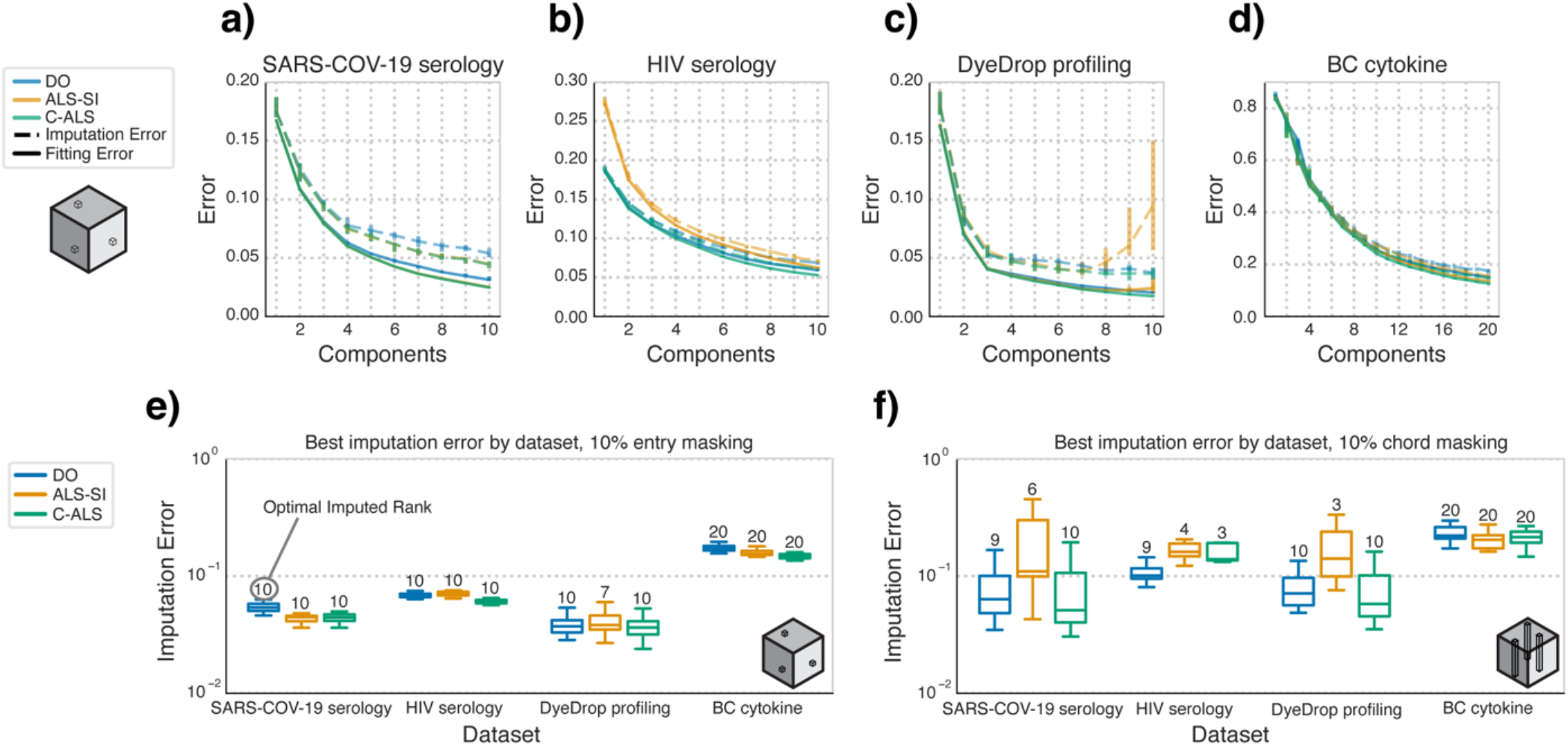
C-ALS outperforms other algorithms in the 10% entrywise masking case. Comparison of the three factorization algorithms in their performance when imputing masked values. **a–d)** Median fitting and imputation error for the **(a)** SARS-COV-2 serology, **(b)** HIV serology, **(c)** DyeDrop profiling, and **(d)** BC cytokine datasets with 10% of the entries masked. **e–f)** Boxplots of the factorization imputation error at the rank with the lowest median imputation error (best imputed rank, labeled atop each bar) per method and grouped by dataset with random entrywise **(e)** and chordwise **(f)** masking. Boxes are defined by corresponding quartile ranges with whiskers denoting the farthest point within 1.5 times the interquartile range.

To summarize this behavior across the choice of different component numbers, we plotted the imputation error for each method at only the optimal imputed rank, determined by selecting the component rank with the lowest median imputed error. With 10% of the entries masked (Fig. 3e), the results recapitulate the earlier results (Figs. 3a–d). With chord-wise masking, C-ALS again maintained consistently similar or lower median imputed error than ALS-SI in all but the BC cytokine dataset (Fig. 3f). Additionally, DO attained lower error than either of the least square methods with the HIV serology dataset, likely due to the large fraction of pre-existing missing values in this dataset. ALS-SI particularly struggled with imputing missing chords, and the best-imputed rank was often lower than that selected with other methods. Under chordwise imputation, the mode along which chords were dropped varied the absolute errors but not how each method compared with each other overall (Table S1).

We then investigated the impact of the amount of artificial masking on the imputation error. We identified the median imputed errors of the optimal imputed rank (Table S2 & S3) for each method, comparing their median imputed and corresponding fitting error for each masking level and type (Fig. 4). The optimal component number for all algorithms at each entrywise masking percentage was generally the highest or second to highest component, except for the least square methods at highest masking percentages (>30%) in the SARS-COV-2 serology and all masking levels in the DyeDrop dataset (Table S1). Still, with entrywise masking, C-ALS obtains the lowest fitting and imputation errors of any algorithm across the datasets even when introducing masking percentages of up to 40% (Fig. 4b-d). The one exception is in the SARS-COV-2 serology dataset, where ALS-SI performs near identically except with marginally lower imputed error at the highest entrywise masking percent (Fig. 4a).

**Fig. 4.**
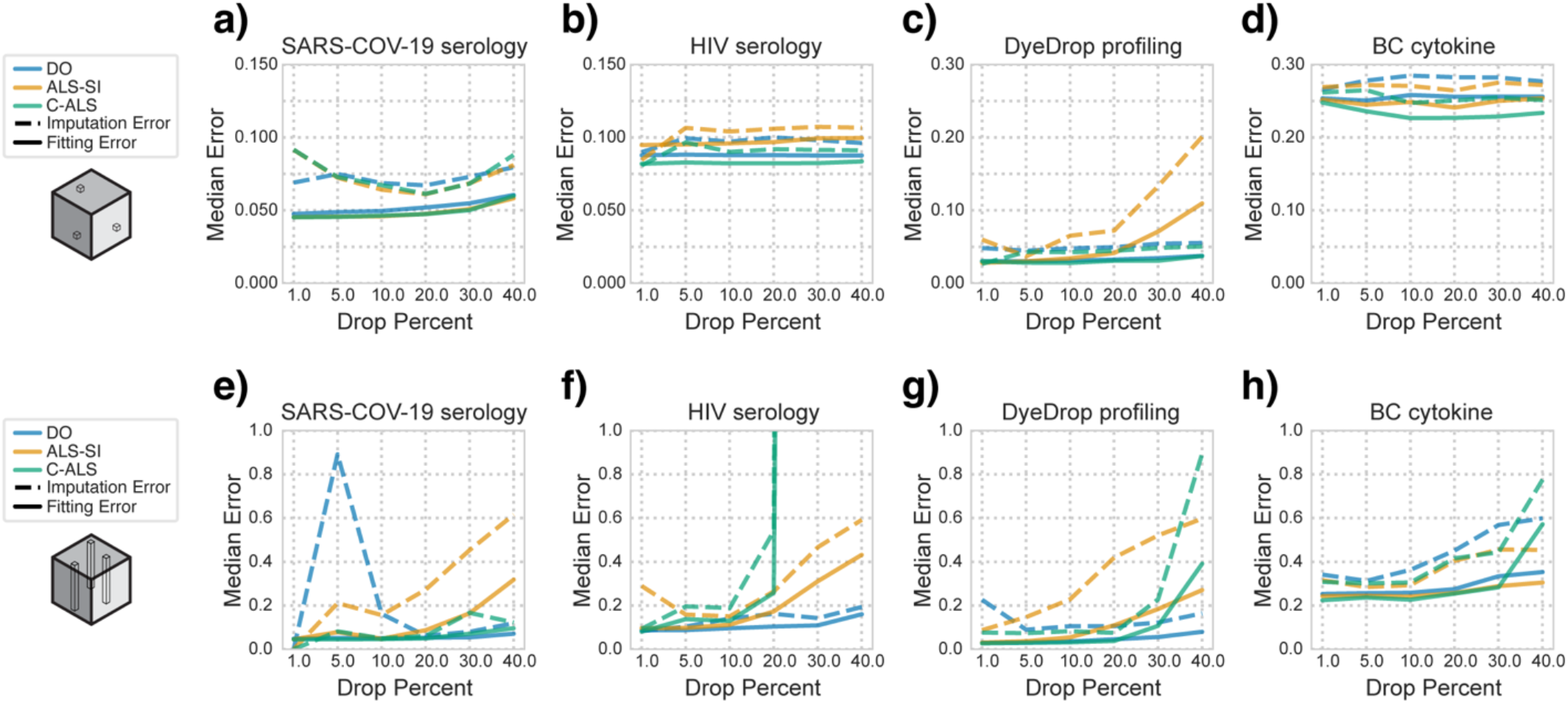
C-ALS outperforms other algorithms with most masking amounts. Median best imputation error at various entrywise **(a–d)** and chordwise **(e–h)** masking percentages as well as the matching fitting error for the **(a**,**e)** SARS-COV-2 serology, **(b**,**f)** HIV serology, **(c**,**g)** DyeDrop profiling, and **(d**,**h)** BC cytokine datasets.

Under chordwise masking conditions, the best algorithm is highly dependent on the dataset. For the SARS-COV-2 serology and DyeDrop profiling, with little or no pre-existing missing values, C-ALS outperforms other algorithms up to 20% chordwise masking (Fig. 4a,c) and at the highest component numbers (Table S1). For imputation of chordwise masking above 20% however, DO outperformed C-ALS in fitting and imputation error (Fig. 4a,c). Similarly, in the HIV serology dataset with a large fraction of pre-existing missing values, DO remained the most stable algorithm with consistently lowest errors at all chordwise masking levels (Fig. 4b). Generally, this mirrors a previous trend where DO is optimal with higher percentages of missing values is present, possibly because underlying patterns are difficult to resolve with such levels of missingness. Meanwhile, for all data but the 4D BC cytokine dataset, ALS-SI was consistently the worst algorithm and peaked in imputation performance at notably lower numbers of components. In the BC cytokine dataset however, C-ALS again outperformed other methods up to 40% masking, after which ALS-SI performed best (Fig. 4d).

### 4.4 Method Behavior and Runtime

We then compared each algorithm on the number of iterations needed to reach convergence. For every dataset (Fig. 5a–d), the number of iterations necessary for DO to fully converge was much higher than the two least squares methods, with or without masking. In the entrywise masking cases (Fig. 5e–f), the errors of C-ALS and ALS-SI decreased at a similar pace over iterations. Though final errors were typically lower for C-ALS, we did not observe meaningful differences between ALS-SI and C-ALS in the convergence rate. For high chordwise masking percentages (Fig. 5g-h), the imputation error of C-ALS increased despite plateauing fitted error. Though C-ALS remained the best method when only considering the fitting error, the steadily rising imputation error indicated overfitting.

**Fig. 5.**
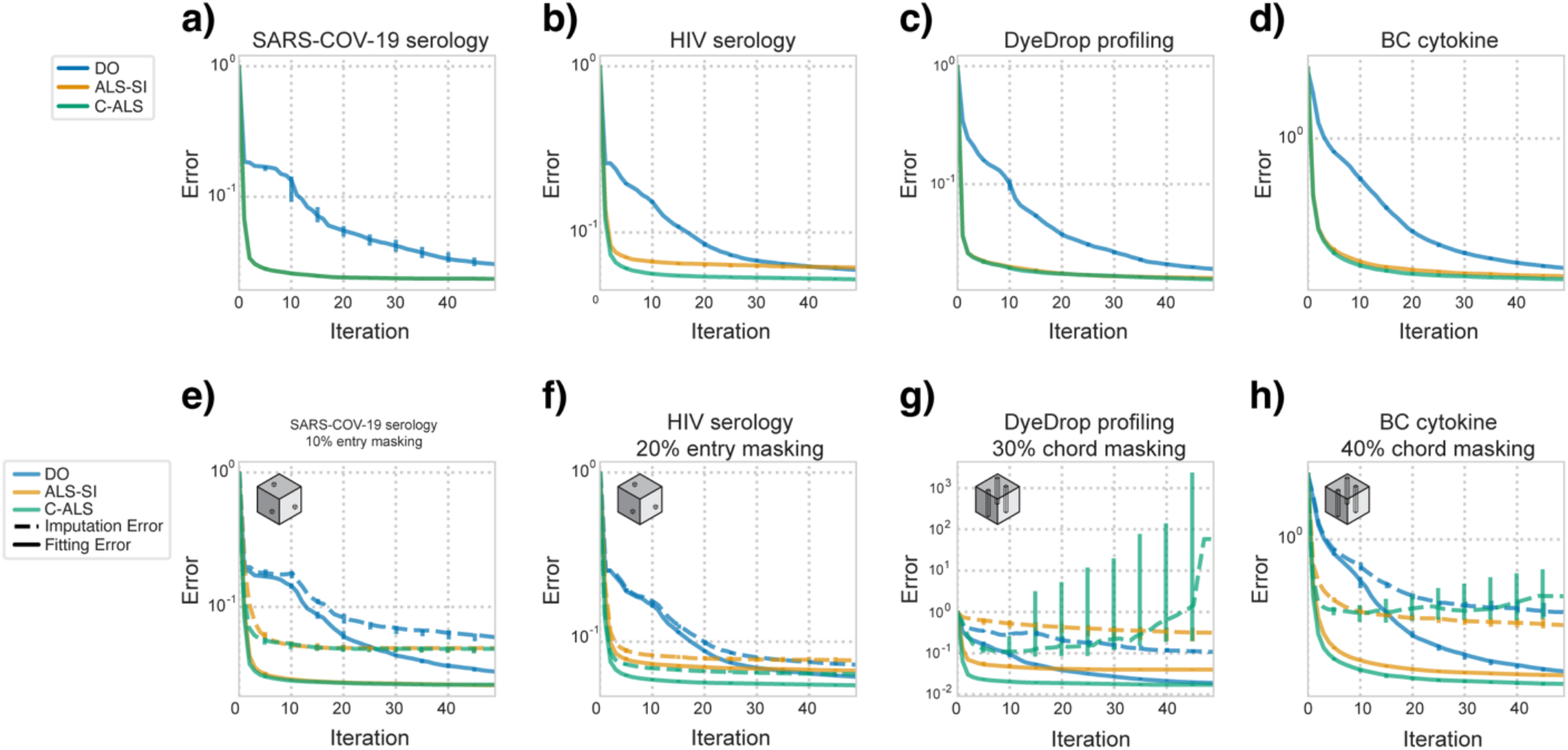
Least squares algorithms converge in fewer iterations than gradient descent and overfitting can be identified. **a–d)** Median fitting error of the indicated dataset over each iteration without masking. **e–h)** Median fitting and imputation error for factorization of the indicated dataset and masking strategy, across iterations, each at the optimal imputed rank.

Although time per iteration was consistent when considering the same masking type on the same dataset, there were slight decreases in time per iteration at higher masking percentages. The one notable exception was that C-ALS had slightly increased time per iteration at intermediate masking percentages for some datasets (Fig. 6b). ALS-SI was consistently the fastest algorithm though at times comparable to DO in both time per iteration. Meanwhile, C-ALS’s time per iteration and time to first iteration relative to other methods varied considerably based on the dataset and masking type (Table S4). In general, C-ALS was noticeably slower than other methods in entry-masking cases (Fig. 6b), but under chordwise masking, C-ALS varied significantly: in the case of the DyeDrop profiling dataset, both methods were faster at low chordwise masking percentages, but these gains diminished as the masking percentage increased (Fig. 6c). The most extreme case difference between the type of masking is observed at 20% entrywise and chordwise masking of the SAR-COV-19 serology dataset: ALS-SI was >10x faster than C-ALS under entrywise masking but <2x faster under chordwise masking (Fig. 6d). C-ALS is at most 4x slower but within 2x of DO’s speed in more than half of the cases (Table S4). In the aforementioned case, C-ALS also had a lower time per iteration than DO (Fig. 6d). Finally, C-ALS used slightly more memory than other methods, though these differences were insignificant relative to those between datasets (Fig. S1).

**Fig. 6.**
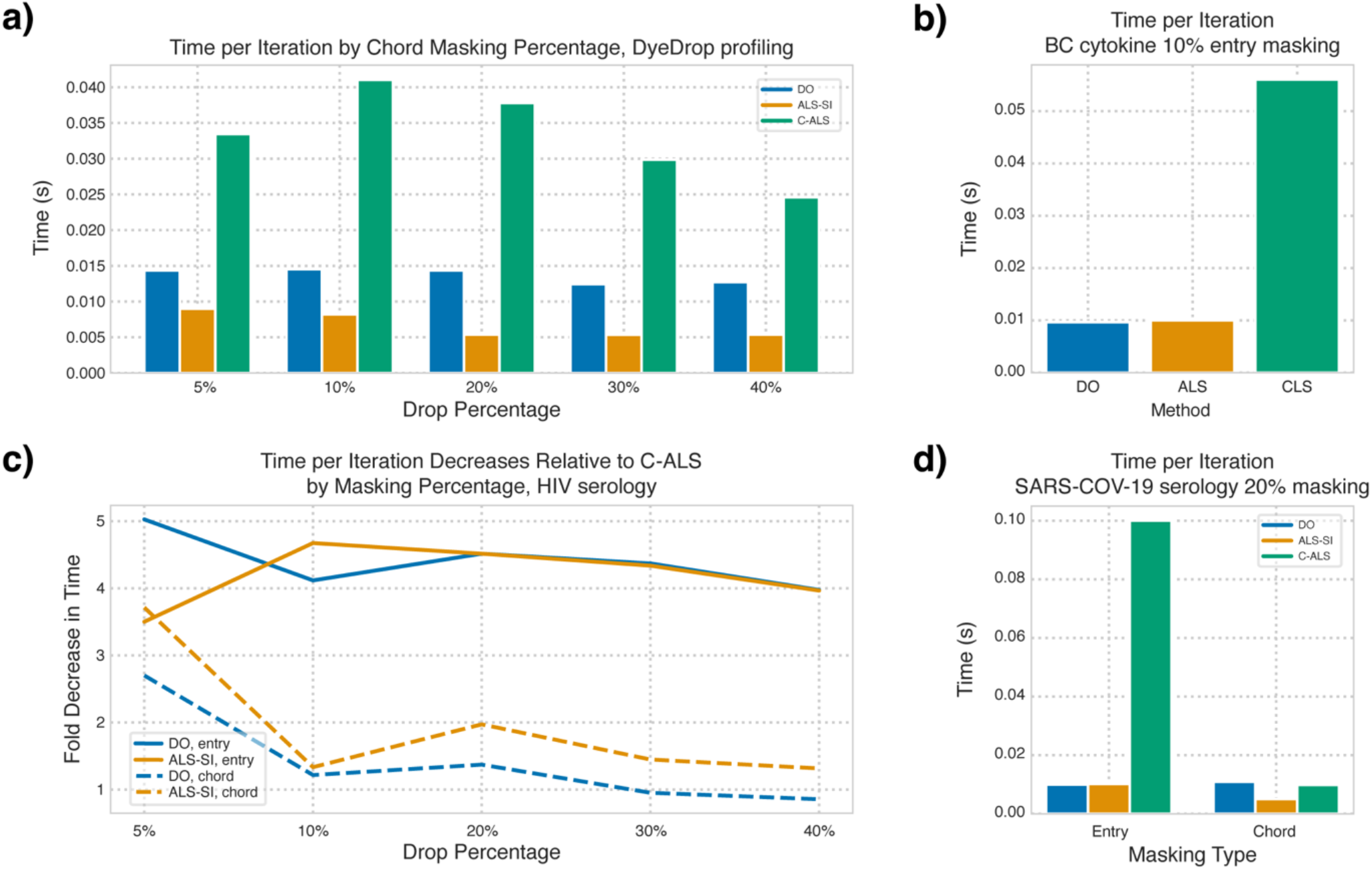
C-ALS is more time consuming per iteration than other methods. **a)** Time per iteration for each method across chordwise masking percentages of the DyeDrop profiling dataset. **b)** Median time per iteration for each method under 10% entrywise masking of the BC cytokine dataset. **c)** Fold decrease in time per iteration for ALS-SI and DO over C-ALS for each method across entrywise and chordwise masking percentages of the DyeDrop profiling dataset. **d)** Median time per iteration for each method under 20% entry- and chord-wise masking of the SARS-COV-19 dataset.

## 5. Conclusions

Our analysis indicates that imputation represents an important metric to consider when selecting a solving algorithm for tensor factorization. As imputed error highlights an aspect of factorization fit not described by other metrics, imputation tests provide insight into determining which fitting algorithm is most appropriate for a dataset. Though running similarly comprehensive imputation testing may be excessive, running imputation on comparable artificial removal patterns and percentages to the original dataset may be useful for identifying the proper solving algorithm, component rank, and identifying issues like overfitting during the factorization process.

To highlight this, when few to no missing values are present, least squares methods consistently achieved better fitting error than DO in all the datasets. Between ALS-SI and C-ALS, C-ALS was generally the better option of the two, but this does not imply however it should be universally applied to datasets. When decomposing datasets with significant amounts of missing values, least squares methods may demonstrate rapidly diverging imputation and fitting errors, suggesting overfitting, that is identifiable at early iterations. In such cases particularly, least squares methods would be a poor choice of fitting algorithm suggesting that imputation error could be an additional convergence criterion useful to consider. Still, while DO appeared to generally be more useful in determining more explanatory patterns from tensor factorization as indicated by comparatively lower imputed error, its reduced accuracy as reflected in the fitting error may become a more pronounced issue for higher rank factorizations. Alternatively, regularization of the C-ALS solving may help to avoid overfitting.

ALS-SI and C-ALS converge in significantly fewer iterations than DO, regardless of the presence or number of missing values, such that when an algorithm is a poor fit for solving the CPD, this behavior may be identified via imputation testing within just a few iterations. While memory is not as notable a concern with any of the given methods, where runtime is concerned, C-ALS must pre-compute multiple linear least squares solutions for even small datasets and is therefore significantly slower per iteration than other methods. As such, C-ALS is necessarily more time-consuming than ALS-SI by a complexity roughly proportional at worst to the number of missing patterns (as observed in intermediate entrywise masking percentages). Additionally, consistent fold-decreases in iteration time of ALS-SI over C-ALS across drop percentages suggest that if time is a concern for very large datasets, sacrificing lower error for a faster solving process by using ALS-SI over C-ALS or DO may be more appropriate on a case-by-case basis. Still, there exists several exceptions to this behavior, and it may be difficult to generalize. Ultimately, the optimal choice of algorithm remains dataset dependent. Considering both imputation and fitting error is key here, as they do not necessarily correspond — suggesting that both error types must be low when solved at a given component rank to be indicative of optimal fit. Still, C-ALS attains these criteria to a greater degree than commonly used methods in the field under a variety of missing value patterns across biological datasets, thus presenting a meaningful improvement to the ALS algorithm for solving tensor factorizations.

## Acknowledgments

None.

## Funding

This work was supported in part by NIH U19-AI172713 to A.S.M.

## Author contributions

A.S.M. and Z.C.T. conceived of the study. E.C.H. and E.H. performed the computational analysis. E.C.H., E.H., and Z.C.T. wrote the first draft. All authors contributed to paper revisions.

## Competing interests

The authors declare that they have no competing interests.

## Data Sharing Plan

All analysis was implemented in Python v3.11 and can be found at https://github.com/meyer-lab/tensor-impute, release 1.0, along with all the experimental data.

## Figures

**Fig. S1.**
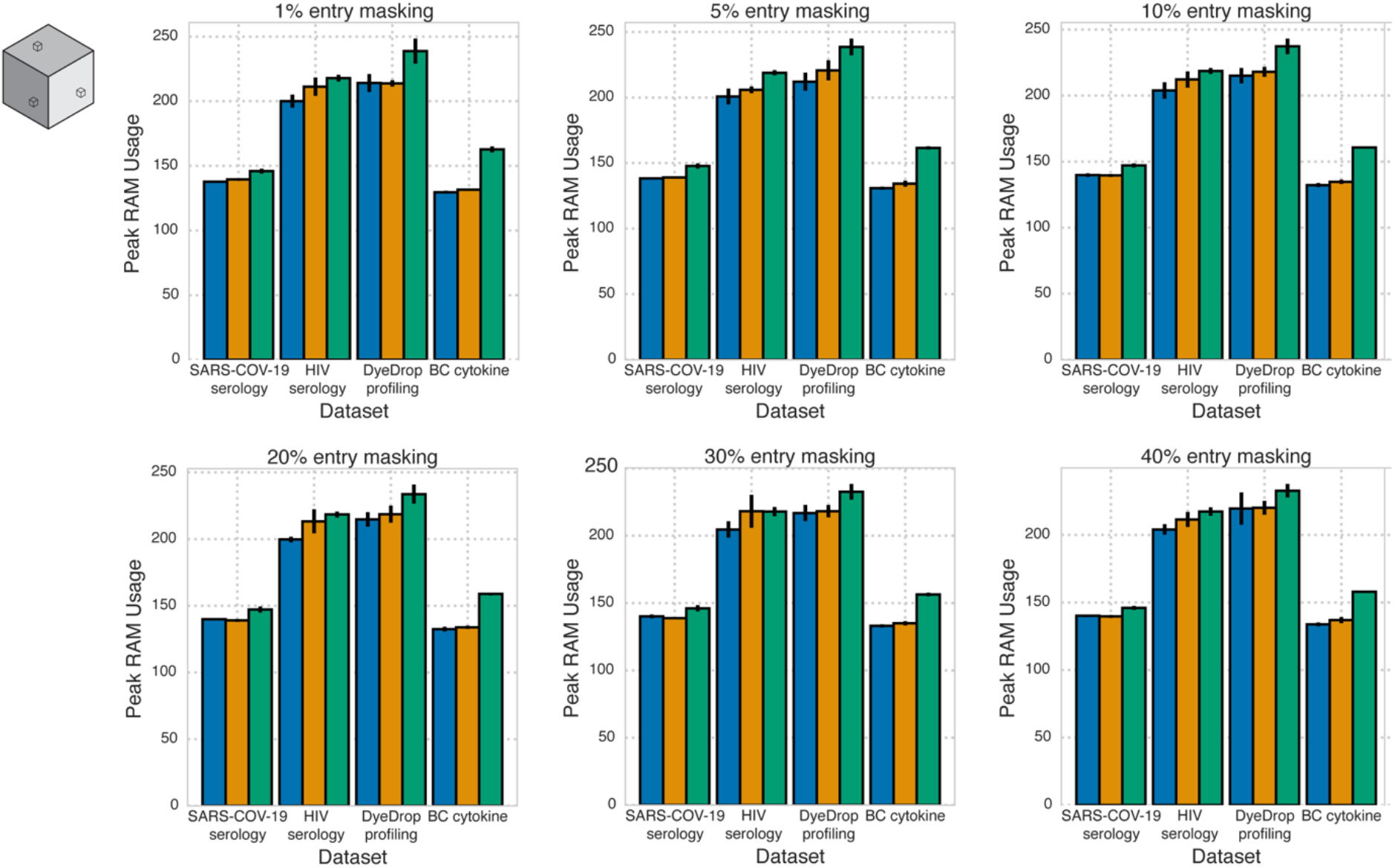
Negligible diZerences exist in memory usage by algorithms across imputation types and extents. Median peak RAM Usage during **(a)** non-missing factorization and at **(b)** 5%, **(c)** 10%, **(d)** 20%, **(e)** 30%, **(f)** 40%, and **(g)** 50% random entrywise masking imputation.

**Table S1. The direction of chordwise masking impacts imputation error but not selection of best solving algorithm.** Median imputation errors at the best imputed rank are provided for chordwise masking along each chord for each solving algorithm at each imputation percentage.

**Table S2. Optimal median imputation errors across entrywise imputation percentages.** Median imputation errors at the optimal imputed rank under each solving algorithm at each entrywise masking percentage.

**Table S3. Optimal median imputation errors across chordwise imputation percentages.** Median imputation errors at the optimal imputed rank for under each solving algorithm at each chordwise masking percentage.

**Table S4. Median time per iteration.** Median seconds per iteration for each dataset for chordwise and entrywise masking combination (rows) for each solving algorithm at each drop percentage (columns). Values displayed are those at the best imputed rank.

## Notes

https://github.com/meyer-lab/tensor-impute

### Competing Interest Statement

The authors have declared no competing interest.

